# Measuring shared responses across subjects using intersubject correlation

**DOI:** 10.1101/600114

**Authors:** Samuel A. Nastase, Valeria Gazzola, Uri Hasson, Christian Keysers

## Abstract

Our capacity to jointly represent information about the world underpins our social experience. By leveraging one individual’s brain activity to model another’s, we can measure shared information across brains—even in dynamic, naturalistic scenarios where an explicit response model may be unobtainable. Introducing experimental manipulations allows us to measure, for example, shared responses between speakers and listeners, or between perception and recall. In this tutorial, we develop the logic of intersubject correlation (ISC) analysis and discuss the family of neuroscientific questions that stem from this approach. We also extend this logic to spatially distributed response patterns and functional network estimation. We provide a thorough and accessible treatment of methodological considerations specific to ISC analysis, and outline best practices.

## Introduction

Traditional methods for fMRI data analysis are not conducive to studying the multidimensional dynamics that characterize social interaction in real-life contexts. Methodological constraints require relatively brief, isolated stimulus events or tasks, accompanied by a predefined model of the expected neural response. Brain areas involved in a particular function are localized by contrasting neural responses to tightly-controlled stimuli varying along a few isolated parameters of experimental interest. As a result, many of the core questions of social and affective neuroscience have proven difficult to study (Zaki and Ochsner, 2011; Hasson and Honey, 2012; Adolphs et al., 2016). For instance, narrative comprehension is triggered by complex situations that unfold over minutes and cannot be captured in brief epochs, while face-to-face social interactions additionally involve a multitude of communication channels such as words, sentences, intonation, facial expressions, and gestures (Hasson et al., 2012). Predicting fluctuations in brain activity during these dynamic, continuous episodes is difficult. Finally, the social and affective symptoms of patients with psychiatric disorders may only be revealed in open, complex situations that cannot be boiled down to experimental paradigms with brief, disjoint events (Klin et al., 2002).

Intersubject correlation (ISC) analysis provides complementary insights to traditional analyses by circumventing the need for a predefined response model and allowing experimenters to measure the consistency of neural responses to complex, naturalistic stimuli across individuals (Hasson et al., 2004, 2010). Beyond simply measuring response reliability, ISC analyses allow us to measure shared content across experimental conditions. By capitalizing on the richness of naturalistic experimental paradigms, ISC has the potential to empower the investigation of social interactions. This tutorial situates ISC among related methods and extends the logic of ISC to spatially distributed response patterns and functional connectivity. We provide illustrative examples of how ISC analyses can be used to address novel questions, and put special emphasis on methodological and interpretational considerations.

### Situating ISC among traditional methods

Traditional analyses of functional magnetic resonance imaging (fMRI) data follow a simple conceptual framework. During experimental design, we generate at least two conditions that differ according to some variable of experimental interest: one may be thought to trigger a particular function while the other is as similar as possible without triggering that function; or the conditions may vary parametrically along a single variable such as retinotopic eccentricity. In fMRI, noise dominates signals at frequencies lower than 0.04 Hz, so when presenting stimuli intended to evoke a particular function, instances of each condition typically range from brief “events” (tens of milliseconds to several seconds) to “blocks” about 20 seconds in duration (Boynton et al., 1996; Chen and Tyler 2008). We assume that neural activity is roughly constant within each instance of a condition, and that a brain region is involved in a function, or tuned to an experimental variable, if its activity increases in response to the condition where the variable of interest is present or increased in magnitude, relative to a condition where the variable of interest is not present (i.e., the control condition) or lesser in magnitude. These are typically referred to as subtraction (Friston et al., 1996) and parametric (Büchel et al., 1998) designs, respectively. These designs lend themselves to generating predictions about the hypothesized time courses of neural activation. The hypothesized time courses serve as predictors in a general linear model (GLM) that quantifies how well the expected time course predicts activity observed in each voxel, thereby localizing the function of interest (Friston et al., 1994b). Fluctuations in brain activity over time within a condition or across repetition are considered noise, while the difference across conditions is the signal. Because each predictor typically comprises multiple trials of the same condition and we assume that neural activity is identical across trials of the same condition, this approach effectively collapses across trials (i.e., trial averaging; Dale and Buckner, 1997). This approach is powerful whenever (*a*) the function can be recruited in short epochs, (*b*) tightly-controlled stimuli can be generated to isolate and manipulate the parameter of experimental interest, and (*c*) we have detailed and exhaustive hypotheses about the time course of relevant brain activity. Cutting-edge modeling efforts (e.g., Huth et al., 2016) suffer from similar constraints. For example, when using word embeddings to predict brain activity during narrative comprehension, each occurrence of a word receives the same embedding regardless of the overarching narrative. In real-life scenarios, where the response to each token changes as a function of an evolving narrative context, such trial-averaging methods will fall short (Ben-Yakov et al., 2012).

ISC analyses provide a complementary, data-driven alternative for identifying brain regions with activity driven by the stimuli or paradigm. The core idea is best illustrated for subjects listening to a spoken story. If multiple subjects listen to the same story, brain regions that are systematically driven by the story will fluctuate synchronously across viewers, while brain regions that do not process the story in the same way across subjects, or are not responsive to the story at all, will not. For example, a voxel in early auditory cortex will consistently track the low-level auditory features of the spoken words across all viewers. The response time course of this voxel will be highly correlated across subjects. On the other hand, a region of the brain that is not entrained by the story (e.g., one involved in low-level visual or motor processing) will not yield a consistent response time course across subjects. Finally, regions that respond to the story in a way that varies temporally to some extent across subjects, for instance because they are involved in emotional reactions to the story that evolve somewhat idiosyncratically from subject to subject, will show intermediate correlations, particularly in the lower frequency range (see Box 1). In summary, correlating brain activity across subjects while they are exposed to a complex stimulus reveals brain areas that process the stimulus in a consistent, time-locked manner. Correlations approaching 1 indicate that the region encodes information about the stimulus and that this information is processed in a stereotyped way across individuals, while correlations approaching 0 reflect regions with idiosyncratic processing or encoding little information about the stimulus.

This logic can be meaningfully applied to specific frequency bands of the signal (Box 1). If we study the processing of features of the soundtrack that fluctuate rapidly, we would look for correlation across viewers in higher frequency ranges. If we study emotional responses that fluctuate slowly, we would look for correlations in slower frequency ranges that also allow for more leeway across viewers in the precise timing of the reaction. Our dependence on the hemodynamic response in fMRI constrains the frequency bands that can be studied with that measurement modality (Box 1). Some of these limitations can be overcome by using other measurement modalities, e.g., ECoG (Mukamel et al., 2005; Honey et al., 2012a), but here we concentrate on fMRI analyses.

Unlike traditional designs where the order of trials may be counterbalanced or randomized across subjects, ISC analysis critically relies on subjects receiving the same time-locked stimulus. Similar to functional connectivity analyses (Friston, 1994a), typical ISC analyses summarize the relatedness of two response time series; however, rather than correlating time series across different voxels within a subject, ISC analyses typically correlate time series across subjects (Figure 1). By computing correlations across subjects rather than across voxels within a subject, ISC analyses are less susceptible to idiosyncratic physiological noise and head motion than functional connectivity analyses (Simony et al., 2016). In another sense, ISC can be understood as specific case of the traditional GLM where the predictor of interest is not generated a priori based on the stimulus or experimental design, but is instead the response time course from the corresponding region in another subject (or the average time course across other subjects). In a traditional GLM, we typically convolve the hypothesized time course of neural activity with a hemodynamic response function (HRF; e.g., Cohen, 1997; Friston, 1998) reflecting the lag and temporal smoothness of the blood-oxygen-level-dependent (BOLD) response. The same HRF is typically used across brain regions, tasks, and subjects, despite evidence for considerable inhomogeneity (Birn et al., 2001; Handwerker et al., 2004). In ISC analyses, there is no need to convolve the hypothesized time course with an HRF, as the hemodynamic responses in one brain are used to predict responses in another brain. Using responses in one brain area to predict responses in the same brain area in another subject mitigates situations in which different brain areas have a systematically different HRFs.

**Fig. 1.**
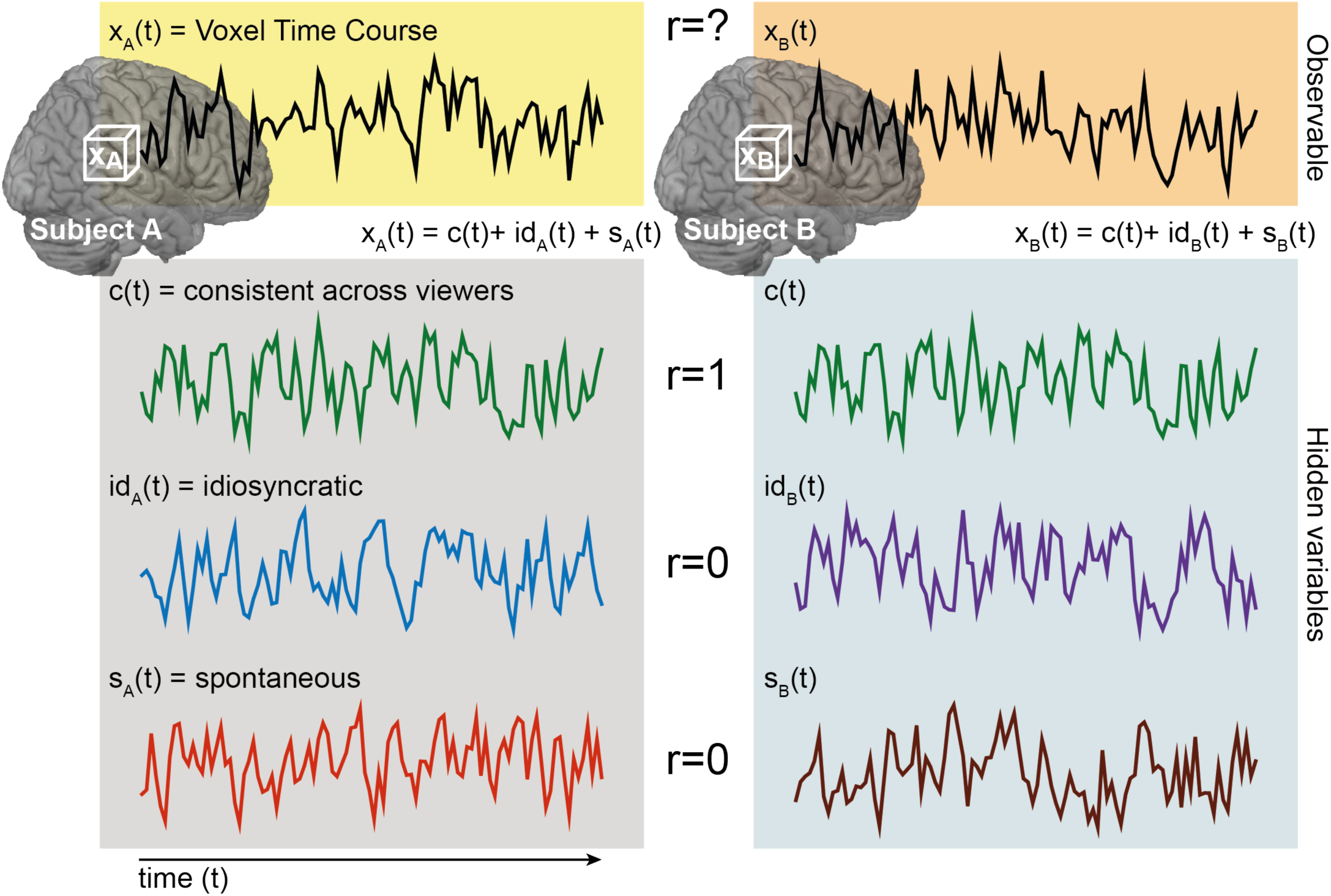
The logic of ISC analysis. The response time course of a specific voxel in a given subject, *x*_*A*_*(t)* can be considered a mixture of three components: a consistent stimulus-evoked component (green), an idiosyncratic stimulus-evoked component (blue), and stimulus-unrelated idiosyncratic or noise component (red). If brain activity is correlated in time across subjects, the green component has a correlation of 1, and the other two components zero. The relative proportion of these components determines the observed ISC.

### Formal definition of ISC

Although we focus on the most commonly-used ISC analysis in this tutorial, this is only one member of a larger family of conceptually related analyses. We first quantitatively consider a typical ISC analysis, then extend this logic to related methods. At the individual-subject level of analysis we can decompose brain activity in a single voxel into several variables (Figure 1). When a given subject *A* listens to a story, the brain activity in a particular voxel over time can be interpreted as a mixture of three signals. The first, which we call *c(t)*, reflects processing that is triggered by the stimulus and is consistent across subjects. For example, brain areas supporting low-level sensory processing closely track stimulus features and respond consistently across individuals. However, stimuli such as stories or movies can also synchronize higher-level brain functions, such as semantic, emotional, and social processing, across subjects in regions beyond sensory cortex (Hasson et al., 2004; Lerner et al., 2011; Thomas et al., 2018). The synchronized component of such higher brain functions is included in *c(t)*. The second variable, which we call *id*_*A*_*(t)*, captures idiosyncratic responses for subject *A* that are nonetheless induced by the stimulus, but with timing and intensity specific to that subject. For example, the same story may be interpreted differently by different subjects if it triggers subject-specific memories or emotions, or the story may evoke similar processes at different times across subjects. The third variable, which we call *ε*_*A*_*(t)*, reflects *s*pontaneous activity unrelated to the stimulus (e.g., thinking about your grocery list during the experiment) and noise (e.g., respiration, head motion). The standardized signal in a voxel *x*_*A*_*(t)* is then a linear combination of these standardized components:

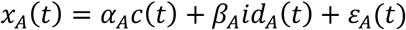

To map all brain regions processing the story, the analysis should quantify how much of the neural activity in each brain region is related to shared and idiosyncratic responses, i.e., *α* + *β* > 0. The larger *α* + *β*, the more the voxel is processing the stimulus. The logic of intersubject correlation is that if a second subject *B* views the same movie, her brain activity will also be a mixture of *c, id*_*B*_ and *ε*_*B*_. By definition, *c(t)* will be perfectly correlated for subjects *A* and *B* (which is why we do not label *c(t)* with a subscript subject variable *A* or *B*), while *id(t)* and *ε(t)* will not be systematically correlated across subjects. By modeling one subject with another subject’s time course, we are effectively filtering out both *id(t)* and *ε(t)*. The actual correlation between the response time course of the two subjects *A* and *B* at voxel *x, r*_*AB*_ = *r*(*x*_*A*_, *x*_*B*_), will thus increase monotonically with *α* (Figure 2), with *r*_*AB*_^2^ ∼ *α*_*A*_ · *α*_*B*_, and with a larger number of subjects, the average 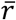 becomes a proxy for the average 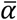. Importantly, intersubject correlation is therefore a tool to detect and quantify shared, stimulus-locked responses, and is insensitive to *id(t)—*a fact that needs to be considered carefully when interpreting results.

**Fig. 2.**
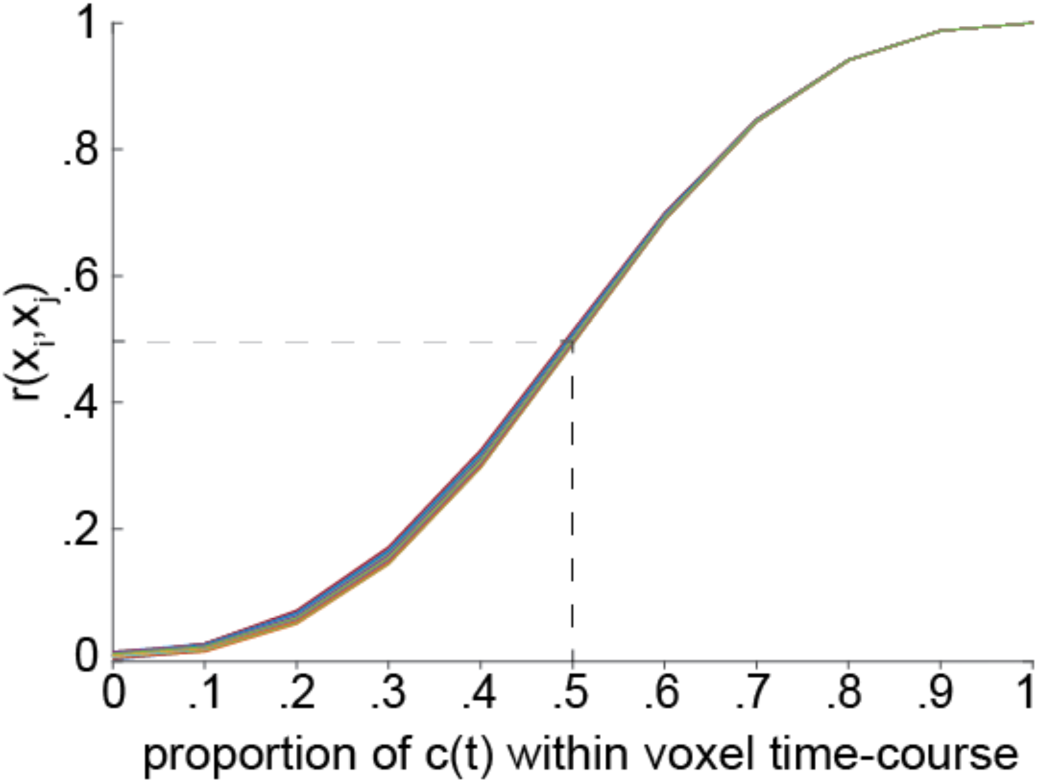
ISC as a proxy for consistent, stimulus-evoked processing. To quantify the relationship between ISC and what proportion of brain activity is consistent, we ran simulations in which response time series were generated for 20 subjects using a mixture of two time series with mean = 0 and *SD* = 1: one was consistent across subjects and one reflected subject-specific noise, with *x(t)* = *α* • *c(t)* + (1 -*α*) • *noise(t)* where *α* is the proportion of consistent signal to noise. The average *r* value over the 20 subjects is shown as a function of *α* (i.e., the proportion of consistent activity). Each line represents one of the 30 simulations. The dashed line illustrates how a case in which 50% of the signal is consistent yields an ISC of 0.5.

Interestingly, although we do not need to know a priori the time course of the consistent, stimulus-evoked component *c(t)* as we must in a conventional the GLM, we can estimate *c(t)* for each voxel from the data, because for large numbers of subjects *N*, 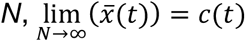; that is, simply averaging the *x(t)* across many subjects provides an estimate *ĉ*(*t*), because the inconsistent components *id(t)* and *ε(t)* will average out to small values close to zero. The main difference between a traditional hypothesis-driven GLM and an ISC analysis is that in the GLM we must have an a priori hypothesis about the time course of activity that is triggered by the experimental design, and then search for regions with this response profile. The stimulus or task is designed so as to generate a specific expected time course. Instead, in ISC analyses we use the shared variance across subjects as a data-driven estimate of *c(t)* and this is done separately for each voxel or region of interest (ROI), allowing each to have a unique time course.

ISC analysis effectively filters out subject-specific signals and reveals voxels with a consistent, stimulus-evoked response time series across subjects. What if, on the other hand, we want to account for idiosyncrasies particular to a given subject? Using the same logic, we can compute correlations within a subject across multiple sessions with the same stimulus. This approach is called “intrasubject correlation” and provides some traction on the reliability of idiosyncratic processes peculiar to individuals (Golland et al., 2007). In the context of intersubject correlation, both *id(t)* and *ε(t)* are uncorrelated across individuals, and the analysis thus isolates *c(t)*. If subject *A* receives the same stimulus multiple times, the correlation between these multiple instances will be sensitive to *id*_*A*_*(t)*, to the extent that the subject-specific processes are stable over multiple exposures. Experience, however, changes how we process stimuli (e.g., Lahav et al., 2007; Engel et al., 2012; Aly et al., 2018). Before measuring intrasubject correlation, one should consider two caveats: first, being exposed repeatedly to the same stimulus leads to habituation (Grill-Spector et al., 2006); second, some idiosyncratic processes are unreliable in their timing, and would thus still fail to register as intrasubject correlation in repeated sessions.

The remainder of this tutorial is divided into two parts. In the first part, we explore how the logic of ISC analyses can be extended to functional network estimation and pattern similarity, and what kinds of scientific questions benefit from these approaches. In the second part, we address practicalities and implementational considerations for designing, analyzing, and interpreting experiments using ISC analyses. Finally, the appendices provide recipes for how to implement these analyses.

## Part I: Extensions and applications

### Temporal intersubject functional correlation

The analyses discussed thus far measure the consistency of responses by computing correlations between homologous brain regions (e.g., the same voxel *x*) across subjects or sessions. However, the logic of ISC can also be used to investigate the functional integration (i.e., connectivity) of diverse brain regions during stimulus processing. We infer that two brain regions are functionally connected if their activity fluctuates in concert (Friston, 1994a). The problem in applying this notion in functional magnetic resonance imaging is that noise in the brain is often shared across voxels. For example, respiration and head motion lead to fluctuations in the BOLD signal across the brain, resulting in spurious inter-voxel correlations that have little to do with concerted neural activity (Power et al., 2012). The logic of ISC offers a way to sidestep these confounds by computing the correlation between the activity of two brain regions *x* and *y* not within a subject, but *across* different individuals—an approach called “intersubject functional correlation” (ISFC) analysis (Simony et al., 2016; Figure 3). Just as *r(*x_*A*_, *x*_*B*_) is a proxy for the amount of information about the stimulus consistently encoded by voxel or brain area *x*, we can extend this reasoning to concerted fluctuations in activity across brain regions *x* and *y* such that *r(x*_*A*_, *y*_*B*_) is a proxy for shared information about the stimulus encoded consistently across these brain regions (Figure 3). That is, ISFC analyses aim to quantify systematic stimulus-evoked communication across brain regions, and can reveal stimulus-related functional networks. ISFC analyses yield a voxel-by-voxel (or ROI-by-ROI) matrix of correlation values for a pair of subjects (or between one subject and the average of others). In practice, computing ISFC yields two asymmetric matrices for *r(x*_*A*_, *y*_*B*_) and *r(x*_*B*_, *y*_*A*_), which are then averaged. The off-diagonal values of this matrix represent functional connectivity between regions, while the diagonal values represent conventional ISCs (each region correlated with itself across subjects). In this sense, the conventional ISC analysis can be understood as a subset of the ISFC analysis (Figure 3). Unlike resting-state functional connectivity analyses, which are intended to measure intrinsic fluctuations (e.g., due to daydreaming) while subjects perform the “rest” task in the scanner, ISFC analyses deliberately filter out idiosyncratic and stimulus-unrelated fluctuations. While traditional functional connectivity analyses yield very similar functional networks whether subjects are at rest or listening to a complex narrative, ISFCs are abolished during rest and very robust during stimulus processing (Simony et al., 2016). Kim and colleagues (2017) have demonstrated that using ISFC analysis to factor out spontaneous activity during a naturalistic vision paradigm yields substantially different functional network solutions compared to rest.

**Fig. 3.**
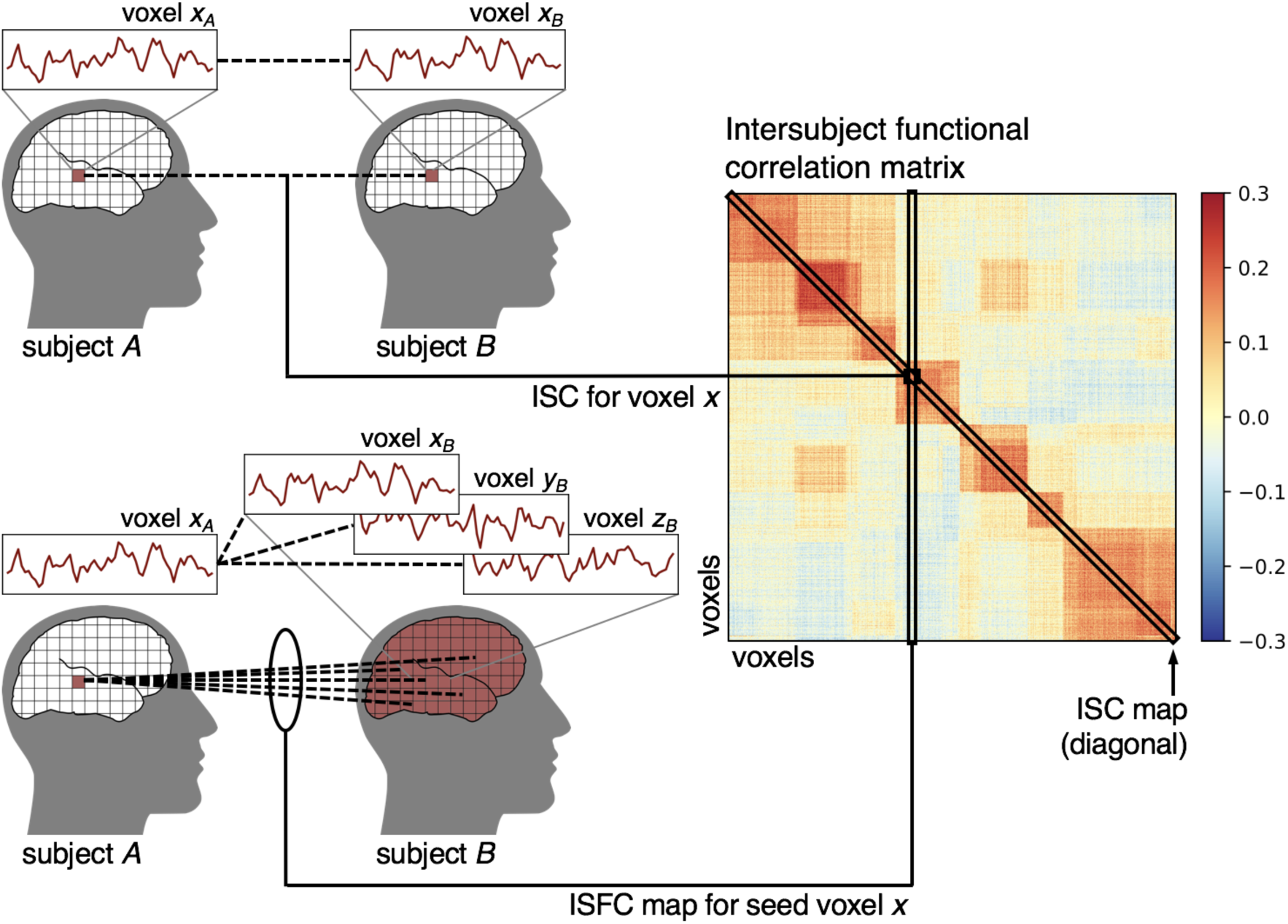
Intersubject functional correlation (ISFC) analysis. Computing intersubject correlations between each voxel and all other voxels yields a voxel-by-voxel intersubject functional correlation matrix. The diagonal values of this matrix reflect the conventional ISC map where correlations are only computed between homologous targets across subjects. A single value on the diagonal corresponds to the intersubject correlation for a given voxel *x*, or *r(x*_*A*_, *x*_*B*_). A single column (or row) of this matrix represents the functional connectivity map for one seed voxel. The off-diagonal values capture all inter-voxel functional correlations, *r(x*_*A*_, *y*_*B*_).

### Spatial intersubject correlation

In addition to computing correlations in response fluctuations over time per voxel or brain area, we can extend the logic of ISCs to multivoxel pattern analysis (Figure 4; Norman et al., 2006; Haxby et al., 2014). In the simplest spatial analogue of ISC (Figure 4A), we compute the correlation between spatially distributed response patterns at a single time point (or the average response pattern across time for a given event) across subjects, thus isolating the shared response pattern *c(s)*, and filtering out idiosyncratic response topographies *id*_*A*_*(s)* and *ε(s)*. This purely spatial approach (referred to as “intersubject pattern correlation”) ignores the temporal evolution of responses, and instead focuses on punctate patterns of activity that are consistent across subjects (J. Chen et al., 2017, Zadbood et al., 2017). Computing the spatial ISC at each time point yields a correlation matrix, analogous to the ISFC matrix (Fig 3), but over time rather than space. In this time-point-by-time-point correlation matrix, the diagonal represents the reliability of the spatial response patterns across subjects at each moment in time, while the off-diagonal values capture whether the same response pattern observed in time *t*_*i*_ is reinstated at time *t*_*j*_. This matrix resembles a time-point representational dissimilarity matrix (RDM) as constructed using representational similarity analysis (RSA), but pairwise dissimilarities are computed across subjects rather than within subjects (Kriegeskorte et al., 2008). Spatially distributed response patterns can be assessed within an ROI or using a searchlight analysis to map local response consistency throughout cortex (Kriegeskorte et al., 2006).

**Fig. 4.**
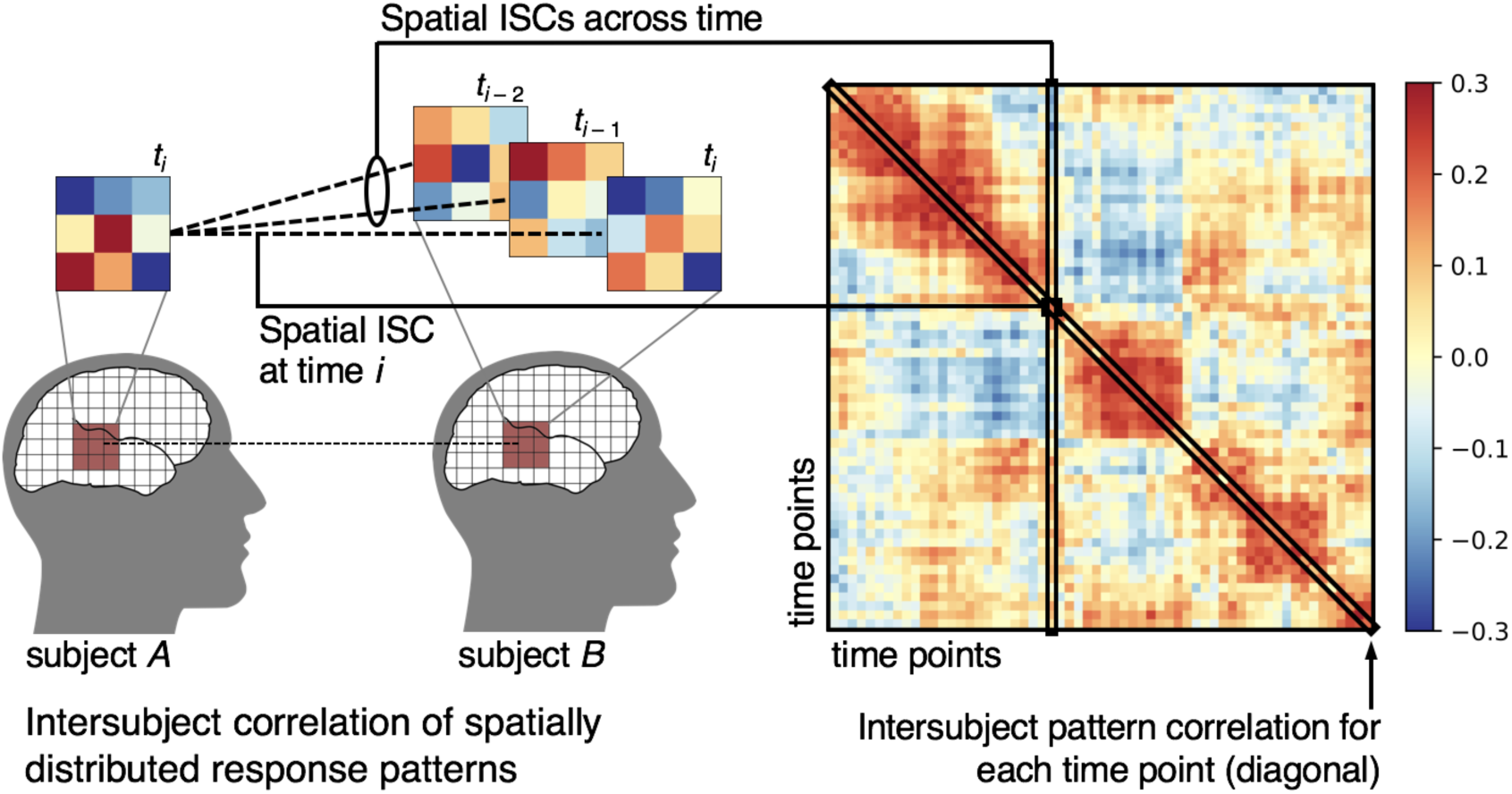
Intersubject correlations of spatially distributed response patterns (“intersubject pattern correlation”). For a given searchlight or ROI, we compute the correlation between response patterns for a single time point *i* (or the averaged response patterns across time points in an event) across subjects. We can also compute intersubject pattern correlations across time points to capture the evolution of response patterns over time (e.g., if a particular pattern recurs at multiple time points). Computing the pairwise intersubject pattern correlations across all time points results in a time-point-by-time-point correlation matrix. The diagonal of this matrix reflects the intersubject pattern correlation at each time point, while the off-diagonal values reflect intersubject pattern correlations across time points.

### Combining temporal and spatial intersubject correlation

Spatial and temporal ISC, while related in many cases, can in principal reveal different, sometimes even complementary, sources of shared responses across subjects. For example, a small region of cortex may yield strong univariate temporal ISCs when response time series are aggregated across voxels, but lack any consistent multivariate variations across space. This would lead to high temporal ISC and low spatial ISC. Conversely, a small patch of cortex may yield consistent spatial response patterns for some time points (or average response patterns across several time points), with inconsistent responses for a given voxel over the entire time series. This would result in high spatial ISC for some time points, and low temporal ISC overall. Even if the aggregate response time series for this region does not yield high temporal ISC, particular time points may have high spatial ISC. To combine the shared signal across space and time we can concatenate spatial response patterns over time, resulting in a multivoxel response trajectory, and assess the intersubject (or intrasubject) consistency of these spatiotemporal response patterns (Feilong et al., 2018; Figure 5). Alternatively, we can apply RSA by computing the pairwise dissimilarities among time points (or conditions) within each subject, then use ISC analysis to quantify the similarity of these time-point RDMs across subjects (Kriegeskorte et al., 2008; Raizada and Connolly, 2012; Charest et al., 2014).

**Fig. 5.**
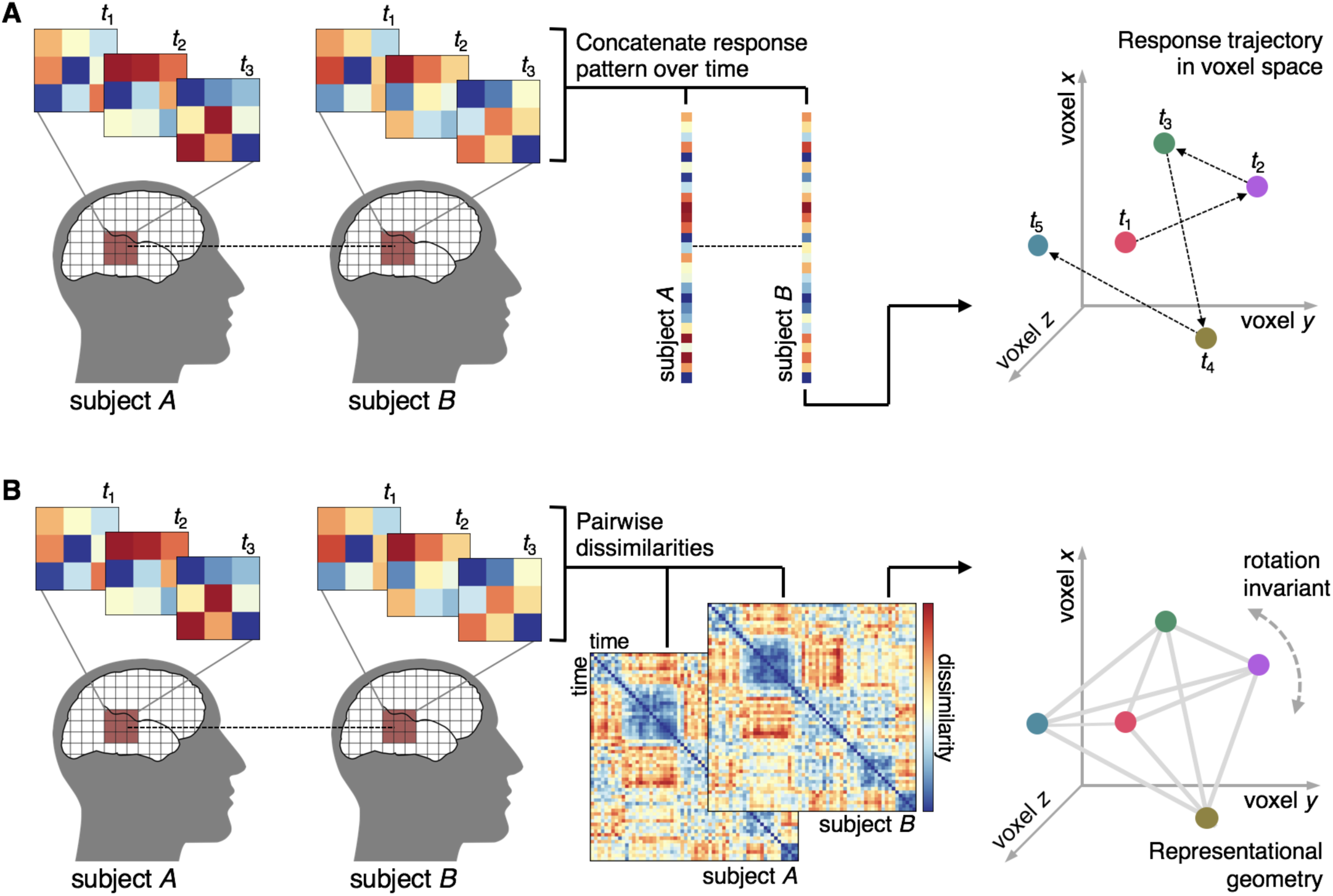
Spatiotemporal intersubject correlations. (A) To quantify the ISC of spatially distributed response patterns over time, we concatenate response patterns for multiple time points for each subject, resulting in a single vector representing a spatially distributed response trajectory, and submit this vector to ISC analysis. The response pattern for each time point can be represented as a vector in a multidimensional space where each dimension corresponds to the response magnitude of a particular voxel; the concatenated spatiotemporal vector is a response trajectory over time in this voxel space. Like the purely spatial approach, this approach requires voxels be functionally aligned across subjects. (B) For a given set of spatially distributed voxels, we can also compute the pairwise dissimilarities between response patterns at each time point to construct a time-point-by-time-point representational dissimilarity matrix (RDM) for each subject. We can then submit the off-diagonal triangle (Ritchie et al., 2017) of this matrix to ISC analysis. Note that representational geometry is invariant to an arbitrary rotation of the response trajectory in voxel space; therefore, computing ISCs using these second-order RDMs abstracts away from each subject’s idiosyncratic voxel space.

## Applications of ISC analysis

What sorts of scientific questions can be addressed using ISC analysis? In the simplest case, computing temporal ISCs across a movie or spoken narrative provides insights into the reliability of stimulus locked neural responses across subjects (Hasson et al., 2010). However, by capitalizing on a shared naturalistic stimulus, ISC analyses can also be used to measure commonalities in stimulus-evoked processing across imaging modalities, such as fMRI, ECoG, EEG, and fNIRS (Mukamel et al., 2008; Liu et al., 2017; Haufe et al., 2018). In this context, ISCs reflect neural signals captured by both measurement modalities. This approach has also been used to explore homologies in neural responses across species (humans and macaques; Mantini et al., 2012). By using interspecies functional correlation analysis in conjunction with a naturalistic visual stimulus, Mantini and colleagues (2012) were able to identify functional homologies across primate species without assuming anatomical correspondence. The same logic can be used to compare neural entrainment to a naturalistic stimulus across populations, such as between autistic patients and controls (Hasson et al., 2009; Salmi et al., 2014) or over the course of development (Cantlon and Li, 2013; Campbell et al., 2015; Petroni et al., 2018; Piazza et al., 2018).

Consider that all the experiments described thus far relied on an identical stimulus. Relaxing this constraint opens the door to a variety of novel questions. To examine how the brain integrates information over time, we can first segment a continuous stimulus, such as a spoken story, at different granularities, such as word or paragraph scales, and present subjects with intact and shuffled versions of the stimulus (Hasson et al., 2008b; Lerner et al., 2011). While responses to the intact stimulus will capture a continuous narrative thread, responses to the shuffled stimulus will not. If shuffled at a very fine scale (e.g., at the level of individual words), ISC will only be high in brain areas with relatively short temporal receptive windows (TRWs), such as early auditory areas. Disrupting the temporal order of a stimulus at an intermediate scale will reveal high ISCs in areas that encode information integrated over longer time periods (i.e., areas with intermediate TRWs), while the fully intact stimulus will yield ISCs across areas with the full range of TRWs. High-level cortical areas encoding features of the narrative that unfold over minutes will only exhibit high ISCs during the intact condition. We can also un-shuffle the brain responses to the shuffled stimulus and compare ISCs across response to the shuffled and intact stimuli. A similar approach has revealed that motor brain region not only process individual observed actions (e.g., grasping) but also contain information about how these actions chain together to achieve meaningful goals (e.g., making breakfast; Thomas et al., 2018).

Qualitatively different stimuli may differentially synchronize brain activity. For example, stimuli varying in emotional content (Nummenmaa et al., 2012), predictability (Dikkers et al., 2014), or audience appeal (Dmochowski et al., 2014) have been shown to yield differential ISCs. Yeshurun and colleagues (2017a) capitalized on the fact that minor stimulus manipulations, such as occasional word substitutions in a spoken story, can radically change the narrative interpretation. Despite surface-level similarity, these stimuli yielded increasingly differentiated responses in higher-level cortical areas.

Even more dramatic stimulus manipulations can be used to isolate systems processing high-level content. For example, Regev and colleagues (2013) presented subjects with spoken and written versions of a narrative. While low-level auditory and visual areas diverged according to presentation modality, particular brain areas yielded high temporal ISCs across modalities indicative of modality-invariant linguistic processing. This approach can also be used to study how complex perceptual stimuli are compressed in memory. In another study, the neural responses of Russian-speaking subjects listening to a story told in Russian were correlated with the responses of English-speaking subjects listening to an English translation of the story (Honey et al., 2012b). This design allows us to identify brain areas which are sensitive to the content of the narrative irrespective of linguistic variations. Chen and colleagues (2017) scanned subjects while they viewed a naturalistic movie stimulus, then instructed subjects to verbally recall events from the movie. They demonstrated that spatially distributed response patterns across subjects in default-mode areas encode event-level representations that are shared across both subjects and across the movie-viewing and verbal recall conditions, indicating a common, high-level representational format for both encoding and retrieval.

Cleverly, instead of directly manipulating the stimulus, we can manipulate attention (Ki et al., 2016, Regev et al., 2018) or narrative context (Yeshurun et al., 2017b). For example, in the report by Regev and colleagues (2018), two distinct and unrelated narratives, one spoken and one written, were presented simultaneously to subjects, and subjects were instructed to orient attention to either the spoken or written story. ISFC analysis was then used to measure how attention routed information from the visual and auditory cortex to higher-order linguistic and extra-linguistic areas (Regev et al., 2018). Yeshurun and colleagues (2017b) manipulated context while presenting two groups of subjects with an identical narrative stimulus. Prior to listening to the stimulus, the groups received brief prompts biasing them to interpret the stimulus according to one of two very different contexts. In high-level cortical areas, within-group ISCs were significantly greater than between-group ISCs, indicating that, despite receiving identical stimuli, the context manipulation resulted in divergent narrative processing. Using an abstract, ambiguous stimulus, Nguyen and colleagues (2019) were able to show that subjects with similar interpretations of the stimulus had similar neural responses.

Finally, one of the most promising applications of ISC analyses, and perhaps the most relevant to the aims of social neuroscience, has been in exploring social interaction across subjects (i.e., brain-to-brain coupling; Hasson et al., 2012; Nummenmaa et al., 2018). For instance, the brain activity of an individual telling a story has been shown to correlate substantially with the brain activity of people listening to that story, and the magnitude of that correlation predicts how well the listener understood the story (Stephens et al., 2010). Similarly, the brain activity of subjects communicating via gestures correlates with that of subjects trying to guess the concept from viewing the gestures (Schippers et al., 2010). Importantly, in these approaches we must consider the fact that there will be variable temporal lags between sender and receiver brains, and the analyses must allow for such shifts. To capture simple delays, we can shift the sender’s voxel time course back and forth in time with respect to the receiver’s response time course, and examine which of these delays leads to the optimal synchrony (cross-correlation analysis; Stephens et al., 2010). Another approach has been to adopt methods that intrinsically accommodate such time shifts, such as Granger causality analysis (Schippers et al., 2010), dynamic time-warping (Silbert et al., 2014), or linear interpolation (Lerner et al., 2014). In the spatial domain, Zadbood and colleagues (2017) extended the results of Chen and colleagues (2017), demonstrating that perception of a naturalistic stimulus, verbal recall, and subsequent narrative comprehension all rely on common, event-level representations encoded in default-mode cortical areas. Finally, recent efforts have used simultaneous “hyperscanning” techniques (Montague et al., 2002; Babiloni & Astolfi, 2014) to extend measurements of brain-to-brain coupling to real-time social interactions (Dumas et al., 2010; Saito et al., 2010; Dumas, 2011; Cui et al., 2012; Jiang et al., 2012; Shilbach et al., 2013), and in some cases going beyond dyads to dynamic group interactions (Jiang et al., 2015; Dikkers et al., 2017).

## Part II: Practical considerations

## Experimental design

A fundamental difference between designing experiments for a traditional GLM analysis and an ISC analysis is that in traditional designs, the main source of signal is the difference in mean amplitude between instances of the conditions (typically collapsed across many trials). Response fluctuations within a condition are considered noise. To increase design efficiency, it is thus best to have many repetitions of the conditions but keep each instance relatively short (less than 20 seconds). This is because noise follows a 1/f distribution, and longer blocks deposit the signal in lower, noisier frequencies. ISC instead uses the fluctuations in activity over time *within* an instance of condition, or over the course of a continuous stimulus, as the signal of interest. Several interrelated factors must be taken into consideration, including the sampling rate, the frequency of neural fluctuations of interest, and the time over which the stimulus conveys meaningful information. Correlations computed over few samples (i.e., time points) are highly unreliable (Fisher, 1921; Bonett and Wright, 2000), so longer epochs are preferred. As a guideline, blocks of duration of at least 30–60 TRs are ideal (Simony et al., 2016); in practice, given a typical sampling rate for fMRI of ∼2-second TRs, stimuli used for ISC analyses often range from ∼1-minute movies (e.g. Thomas et al., 2018), ∼5-minute narratives (e.g., Lerner et al., 2011), to feature-length films over 1 hour long (e.g., Hasson et al., 2004; Haxby et al., 2011). Note that while some neuroimaging modalities have much higher temporal resolution than fMRI (e.g., ECoG), the neural signals most reliably shared across individuals may nonetheless fluctuate relatively slowly (Honey et al., 2012a). To understand the minimal duration of epochs, it is thus important to consider the frequency of the signals one is interested in measuring, and how different cortical areas may be sensitive to information evolving over different time scales. For example, to capture consistent, stimulus-evoked processing in prefrontal regions, we need to present subjects with a coherent stimulus which unfolds over at least several minutes (Hasson et al., 2008b; Jääskeläinen et al, 2008; Lerner et al., 2011). Note that the window size used to compute ISC and the coherence of the stimulus are independent parameters. We can use a relatively brief sliding window of 30 TRs to compute dynamic ISCs during a coherent 1-hour movie to assess how signal in higher-order brain areas fluctuates over time as the movie unfolds. However, using a 30 TR sliding window and scrambling the movie at the scale 10-second segments will not capture reliable responses in these higher-order areas. Thus, to capture responses with long processing timescales, we advise using coherent stimuli which unfold over minutes (Hasson et al., 2015). When using sliding-window ISCs to measure fluctuations in synchrony, there is a tradeoff between the temporal resolution at which fluctuations and the reliability of the ISC estimate when determining the width of the window. Related metrics such as intersubject phase synchrony that capture instantaneous, time-varying synchronization may provide additional insights into dynamic intersubject coupling (Glerean et al., 2012).

ISC analyses—because they do not require an explicit model of the task or stimulus—are particularly useful for naturalistic experimental paradigms, where constructing such a model may be prohibitively difficult. Relative to traditional fMRI experiments that typically use highly-controlled stimuli, naturalistic stimuli are more ecologically valid (Zaki and Ochsner, 2011; Hasson and Honey, 2012; Adolphs et al., 2016; Hamilton and Huth, 2018), convey rich perceptual and semantic information (Bartels and Zeki, 2004; Huth et al., 2012, 2016), and more fully sample neural representational space (Haxby et al., 2011, 2014). Recent work (Vanderwal et al., 2015) also suggests that naturalistic stimuli may improve subject compliance (in terms of wakefulness and head motion relative to, e.g., rest), which is particularly important when scanning patient populations and children. As mentioned previously, different stimuli will variably synchronize different brain systems; for example, engaging, Hollywood-style movies may yield greater, more widespread ISCs than real-life, unedited videos (Hasson et al., 2010; Cohen et al., 2017).

Conventional ISC analyses critically depend on temporal similarity across subjects. It is therefore essential to have the scanner hardware trigger the computer controlling the paradigm to start the stimulus at the same time across subjects; logging all trigger pulses from the scanner will ensure that stimuli are presented at the same moment, relative to each acquired volume, in all subjects. If the design is divided into multiple epochs, the order of the epochs can be randomized across subjects during the data acquisition, and rearranged to a common order prior to ISC calculation (assuming there is no narrative structure across epochs).

In theory, block and event-related designs can be analyzed using an ISC approach if stimuli were presented with exactly the same timing for all subjects (Hejnar et al., 2007; Pajula et al., 2012). In that case, the entire functional time course can be correlated across subjects (cf. Bordier and Macaluso, 2015). However, if data from both rest and stimulus periods are correlated across subjects as a single time series, increases and decreases in the BOLD signal will be largely driven by the onset and offset of each event, yielding ISC maps resembling the activation maps of a traditional GLM analysis. Experiments designed for a traditional GLM analyses often do not use identical trial orders across subjects to avoid confounding order effects, and may have variable event durations due to subject-specific behavioral responses. To concentrate on the specific processing during a task, it is essential to splice the data to exclude rest, onsets, offsets, and compute ISC only during the task. To exclude onsets and offsets entirely, we recommend removing the first 10 seconds of data of each epoch and only considering data up to the end of the stimulation epoch. Note, however that these transients may last considerably longer than 10 seconds, and may vary across subjects, stimuli, and brain areas. Visually inspecting the response time series in representative ROIs may be informative for gauging the duration of transients. This need for trimming further motivates designs with relatively long epochs. After splicing and trimming, time series from each block of a given condition should then be standardized (*z*-scored) prior to concatenating segments to avoid introducing large signal changes at the joints (see Appendix A for details on preprocessing data). If blocks comprise different conditions, data from all the blocks of a given condition can be concatenated to generate an ISC estimate per condition to be compared at the second level across conditions.

## Computing ISC and statistical inference

Like most fMRI analyses, conventional ISC analyses follow the historical approach of dividing the statistical analysis into two stages: individual-subject (first-level) and group (second-level) analyses. At the first level, we assess the similarity of brain activity across different subjects, while at the second level, we assess whether this level of similarity is significantly greater than zero or significantly different across groups or conditions.

At the individual-subject level of analysis, we use Pearson correlation to measure the statistical association between the response time course for one subject and other subjects at each voxel or ROI. The Pearson correlation coefficient measures the linear association or dependence between two continuous variables. Note that Pearson correlation is scale-invariant; that is, Pearson correlation implicitly mean-centers and scales the input variables to unit variance (i.e., *z*-scoring). These properties of the Pearson correlation coefficient also apply to spatial approaches to ISC (i.e., effectively mean-centering regional response magnitudes; Misaki et al., 2010). There are two commonly-used approaches for computing ISCs at the individual-subject level:

*Pairwise approach*: In this approach, each subject is correlated with every other subject, leading to *N*(*N –* 1)*/*2 *r*_*AB*_ values, where *r*_*AB*_^2^∼*α*_*A*_ · *α*_*B*_ and the average of these r-values, 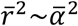. The resulting correlation values are typically represented as a symmetric subject-by-subject correlation matrix where each cell of the upper (or lower) triangle reflects the ISC between a pair of subjects.

*Leave-one-out approach:* The other approach leverages the fact that *c(t)* can be approximated by averaging the response time course *x(t)* over subjects. For every subject *A*, we can then approximate *c(t)* by averaging over all other subjects (i.e., excluding subject *A*), and get an approximate *α* for each subject using 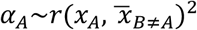. Using this approach, we obtain higher *r* values than using the pairwise approach, because *r* in the pairwise approach is a function of *α* while in the leave-one-out approach *r* is a function of 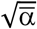; if 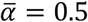, the pairwise approach will lead to *r* values around 0.5, while the leave-one-out approach will have values around 0.71. We obtain *N* estimates (one per subject), instead of *N*(*N-*1)/2 as in the pairwise approach.

At the second level, we draw inferences about shared activity at the population level. Performing group-level statistical tests for population inference in the context of ISC analyses is surprisingly complex (G. Chen et al., 2016, 2017b), and we will point to the core problems below. In general, for one-sample tests—i.e., testing whether the mean ISC is significantly greater than zero—we recommend using either time series randomization (circular time-shift or phase randomization) or the bootstrap hypothesis test. For two-sample tests, we recommend using a permutation test that randomizes condition or group assignments, or, for those familiar with traditional data analysis packages like AFNI, FSL, or SPM, the conventional two–sample *t*-test.

In traditional GLM analyses, first-level models are constructed independently per subject, and the resulting parameter estimates (e.g., regression coefficients or contrasts) are submitted to a group-level analysis where subject is modeled as a random effect. In ISC analyses, however, each subject typically contributes to the first-level model for every other subject. This means that, in both the pairwise approach, and to a lesser extent the leave-one-out approach, the resulting correlation coefficients are not statistically independent samples and therefore violate the assumptions of common parametric tests (e.g., *t*-test, ANOVA). For example, in the pairwise approach, each subject contributes to (*N* – 1)/2 pairs, leading to highly interdependent correlation values and artificially inflated degrees of freedom. A further problem is that fMRI data follow a power law, and such data can generate spurious correlations (Schaworonkow et al., 2015). Parametric tests, such as the one-sample *t*-test, should thus never be used test the significance of pairwise ISCs. For the leave-one-out approach, especially when comparing ISCs across two conditions or groups, these problems are somewhat attenuated: the issues related to power laws and non-independence are likely to influence both conditions or groups similarly. Accordingly, when comparing two conditions using the leave-one-out approach, two-sample *t*-tests yield robust results that are very similar to the nonparametric tests described below (e.g., Thomas et al., 2018). People with limited programming experience may opt for this approach as it can be easily integrated into traditional analysis packages such as AFNI, FSL, or SPM (Appendices A and B). Further validation of this approach is however ongoing.

In general, there have been two main approaches for statistical evaluation in the literature. These approaches are implemented in either the freely available ISC Toolbox (Kauppi et al., 2014) or the Brain Imaging Analysis Kit (BrainIAK, https://brainiak.org; see Appendices C and D for basic usage of these toolkits; see). In the context of conventional ISC analysis, the first approach assumes that if response time series are correlated across subjects due to time-locked shared neural responses, shifting one of the time series back or forth by a random interval should disrupt the temporal alignment and attenuate the correlation (while still preserving the temporal autocorrelation structure of the response time series; Kauppi et al., 2010; 2014). In this resampling approach, each time series is randomly shifted many times (e.g., 10,000 times), and 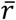 is calculated each time to generate a null distribution of 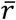 values. The actual value of 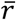 obtained from the original data is then ranked among the time-shifted null distribution, resulting in a *p*-value. The closely related phase-randomization approach (e.g., Lerner et al., 2011) proceeds by applying a fast Fourier transform to the time series, randomizing the phase of each Fourier component, then inverting the Fourier transformation, thus preserving the power spectrum of the signal but disrupting the temporal alignment. Phase randomization is performed at each iteration of the resampling procedure, prior to computing ISC, and the resulting 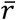 values are aggregated into a null distribution. Both of these nonparametric approaches apply randomization at the level of the time series, and require ISCs to be recomputed at every permutation, making them computationally intensive.

The second main approach operates directly on the ISC values (e.g., *r*_*AB*_ in the pairwise approach) for group-level inference and includes both nonparametric and parametric procedures (G. Chen et al., 2016, 2017b). Chen and colleagues (2016) have suggested that the above approach based on randomized temporal offsets may result in inflated false positive rates (FPR). To account for this, they advocate for two nonparametric approaches that better control the FPR. For one-sample tests using the pairwise approach (where *H*_*0*_: 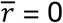), they propose a subject-level bootstrap hypothesis testing procedure. At each iteration of the bootstrap, *N* subjects are randomly sampled with replacement, the ISCs for the resulting sample of subjects is retrieved, and then the test statistic is computed across these pairs. Because this is a nonparametric test, we compute the median ISC rather than the mean (Chen et al., 2016). Repeating this procedure many times (e.g., 10,000 times) yields a bootstrap distribution. Note that constructing a correlation matrix while sampling with replacement will yield off-diagonal 1s when computing ISC for the same subject sampled more than once. We recommend excluding these values when computing the median ISC for each bootstrap sample (Nili et al., 2014). To test the hypothesis, the null distribution should be normalized by subtracting the actual median correlation from each bootstrap median, and the actual median correlation is ranked against this distribution (Hall and Wilson, 1991). For two-sample tests, Chen and colleagues (2016) recommend using a subject-level permutation test to control FPR. In this procedure, group assignments are randomly permuted at each iteration, effectively exchanging entire rows/columns of the pairwise ISC matrix. Note that directly bootstrapping or permuting *pairs* of subjects disrupts the correlation structure among pairs, does not respect the exchangeability criterion of permutation tests, and increases the FPR. Finally, G. Chen and colleagues (2017) propose a parametric linear mixed-effects modeling procedure with crossed random effects indicating which subjects contribute to each pair so as to account for the correlation structure among pairs. This approach has greater flexibility (e.g., can accommodate covariates) and potentially lower computational cost than nonparametric tests, but relies on stronger assumptions. Because these approaches operate on ISC values rather than the response time series, they are also applicable to spatial ISC methods.

Finally, note that Pearson correlation coefficients used during ISC analyses should be Gaussianized via the Fisher *z*-transformation (inverse hyperbolic tangent function *arctanh*) prior to averaging, as simple averaging will tend toward a downward bias (Fisher, 1915; Silver and Dunlap, 1987; Chen et al., 2016). Fisher *z*-transformation is important for any parametric statistical test; in the case of nonparametric methods, the median Pearson correlation should be preferred to the mean (Chen et al., 2016). If you opt to report average Pearson correlations (e.g., when plotting ISC maps), they should be Fisher-transformed prior to averaging, and then the average should be inverse Fisher-transformed.

These statistical tests are often performed independently for every voxel in the brain, introducing a pernicious multiple testing problem (Nichols, 2012). A common approach is to control the false discovery rate (FDR), which sets the proportion of false positives among detections to a low value such as .05 (Benjamini and Hochberg, 1995; Benjamini and Yekutieli, 2001; Genovese et al., 2002). Controlling FDR in this way ignores the spatial structure of ISC values across voxels and lacks any spatial specificity; we cannot conclude that any particular voxel in an FDR-corrected ISC map is significant, only that no more than 5% of the detected voxels are false positives (Poldrack et al., 2011, pp. 121–123). A common alternative is to control the family-wise error rate (FWER) using cluster-extent based thresholding (Nichols and Hayasaka, 2003; Woo et al., 2014), which takes the spatial contiguity of brain signals into account. In this approach, we set a cluster-forming threshold (e.g., *p* = .001; cf. Smith and Nichols, 2009) then assess the significance of clusters of voxels that survive this threshold by modeling the distribution of clusters occurring by chance using either random field theory (RFT; Worsley et al., 1992), Monte Carlo simulation (Forman et al., 1995), or permutations (Nichols and Holmes, 2001; Eklund et al., 2016). Note that cluster-wise inference methods suffer from a similar spatial specificity problem: we cannot conclude that any particular voxel or peak within a cluster is significant, only that the cluster as a whole is significant. In the permutation-based approaches described above, instead of constructing a null distribution of 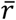 values for each voxel, we can aggregate the maximum 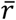 value across all tested voxels at each iteration of the permutation test, resulting in a null distribution of maximum 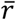 values. We can reject the null hypothesis for any voxel where the observed 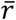 value exceeds our threshold for statistical significance based on this null distribution of maximal 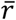 values (e.g., observed 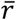 values in the top 5% of the null distribution), strongly controlling the FWER (Nichols and Holmes, 2001; Simony et al., 2016). For simplicity, we suggest controlling FDR in keeping with the precedent in the field, while bearing the above limitations in mind.

## Interpreting ISC results

Traditional fMRI analyses localize task-related increases or decreases in BOLD activity and results are typically described as “region *x* is activated by task *T*.” ISC results are interpreted differently. If a region *x* shows significant ISC (or greater ISC for one task or group than another), we do not conclude that region *x* is activated by the stimulus, but rather we infer that region *x* encodes information about the stimulus that is consistent across individuals. If there is significant ISC, the response time course in one subject’s brain predicts that in another. Similarly, significant ISFC indicates that the response time course in one brain area predicts that in another brain area across subjects. Because the only thing in common between subjects across time is the experimental paradigm, this cross-subject relationship must be mediated by the paradigm—there is thus mutual information between the stimulus and the neural response. This is borne out by the demonstration that one can reconstruct with high fidelity the sound envelope of a movie simply by looking at the shared brain activity in early auditory cortex (Honey et al., 2012a). Importantly, positive ISC in a brain region can be induced by both consistent increases and decreases in brain activity across subjects, and thus should not be interpreted as increased activation across subjects. Given that the nervous system often encodes information using reductions of firing rates (Dacey, 2000), and that reductions in BOLD activity have been related to important brain functions (Anticevic et al., 2012), we feel that moving away from activation can be fruitful. In short, significant ISC reveals that there is a relationship between brain activity and the stimulus, but not the nature of that relationship.

A simple way to gain qualitative insights about what is encoded in a brain region with significant ISC is to explore the average signal across subjects in that region. As mentioned earlier, the average time course of activity across subjects reflects *c(t)* and is representative of the systematic response to the stimulus. Moreover, periods in which the stimulus failed to recruit that brain region will average to zero, while periods in which the stimulus caused consistent activation or deactivation will exhibit significant positive or negative deflections. Assuming a hemodynamic delay of ∼5 s, we can inspect the stimulus for systematic features occurring prior to these peaks (Hasson et al., 2004). To expand this approach to the entire brain, at least two related possibilities exist. First, we can calculate the average 4D brain activity (i.e., the 3D volume across time), and then submit this to independent component analysis (ICA) to summarize the varying time series throughout the brain. The time course of each IC can then be examined and related to the paradigm (Lahnakoski et al., 2012). Second, if the ISC analysis identifies a large number of regions surviving a particular statistical criterion, one can submit the average time courses throughout the brain to a clustering algorithm to functionally parcellate the cortex or identify ROIs with similar time series (Kauppi et al., 2010b, 2017; Thomas et al., 2018). Alternatively, ISC can be computed using a sliding-window approach in order to identify epochs in which ISC was highest, which can then be related to the stimulus (window sizes in the literature range from 10 to 60 TRs; Nummenmaa et al., 2012; Simony et al., 2016).

It is important to quantify the selectivity of neural responses within and across regions. On the one hand, we observed that different brain regions along the processing hierarchy respond differently, resulting in high within-region correlation across subjects and low inter-regional correlations (Hasson et al., 2010, Hasson et al., 2015). If different brain areas have unique response profiles and the resulting region-by-region correlation matrix is meaningfully structured, this suggests that the observed ISCs are not simply due to non-specific or non-neuronal variables like arousal and stimulus-correlated head motion. However, we also observed that intersubject correlation across brain areas belonging to the same functional network (e.g., different areas within the default-mode network) tend to have stronger stimulus-locked covarying activity than areas sampled from different networks. This discovery motivated the development of ISFC analysis (Simony et al., 2016).

In addition to relating ISCs to the stimulus, we can also relate ISCs to behavioral measures. For example, Hasson and colleagues (2008a) used a subsequent memory paradigm to index which events of a movie viewed in the scanner would be remembered three days later for each subject. A voxelwise pairwise ISC analysis revealed brain areas (e.g., parahippocampal gyrus, temporoparietal junction) where ISC was greater for events remembered by both subjects. In addition to item-or event-level episodic recall, aggregate comprehension scores can be related to the spatial extent or magnitude of ISCs (e.g., Stephens et al., 2010).

Finally, as a measure of response reliability, inter-and intrasubject correlations can play an important role in setting an upper bound for the stimulus-related information we can hope to extract from a response time course and can be used to estimate a “noise ceiling” to which models can be compared (Huth et al., 2016; Nili et al., 2014). At a procedural level, ISC analyses can be used as a method for excluding outlier subjects or for feature selection prior to subsequent analysis; e.g., restricting an analysis to only ROIs with high consistency across subjects or ROIs with particular processing timescale (e.g., Yeshurun et al., 2017a; cf. Kriegeskorte et al., 2009).

## Limitations

ISC analyses allow us to leverage more complex stimuli and paradigms, but also have limitations that need to be considered carefully when designing experiments and interpreting results. Critically, ISC analyses require the fluctuations of brain activity to roughly correspond across individuals in both time and space (Box 1). In the temporal domain, methods such as dynamic time-warping can accommodate temporal mismatch and can in part alleviate this limitation (Lerner et al., 2014; Silbert et al., 2014). In the same vein, typical ISC analyses only measure linear associations in activity across subjects (cf. Glerean et al., 2012) and ISFC analyses cannot capture nonlinear transformations occurring between brain areas (Anzellotti and Coutanche, 2018). In the spatial domain, slight misalignments in functional–anatomical correspondence across individuals can dramatically reduce observed ISC values if brain activity is measured at a high spatial resolution without smoothing. While coarse-grained spatial response topographies may be preserved across subjects (e.g., J. Chen et al., 2017), functional alignment algorithms such as hyperalignment resolve idiosyncrasies in fine-grained functional topographies across subjects and can considerably improve both spatial and temporal ISCs (Haxby et al., 2011; Chen et al., 2015; Guntupalli et al., 2016; J. Chen et al., 2017; Feilong et al., 2018)

Second, BOLD activity is strongly affected by respiration, and certain stimuli are known to entrain respiration (Codrons et al., 2014). Although ISC analyses filter out idiosyncratic noise, synchronized stimulus-related respiratory and motion artifacts may contribute to ISCs. Regressing out BOLD signals from the CSF and white matter during preprocessing may provide some protection against such stimulus-correlated noise (at the cost of reducing sensitivity to stimulus-evoked effects of interest). Consult Simony and colleagues (2016) for a more detailed examination on the possible contribution of physiological measurements to ISC analysis.

Finally, our discussion has largely been limited to ISC analyses as they have historically developed in the fMRI community. Closely related analyses have in fact expanded outside the context of neuroimaging; for example, to measuring intersubject synchrony of pupil dilation (Kang and Wheatley, 2017) and gaze direction (Hasson et al., 2008b; Shepherd et al., 2010; Wang et al., 2012). On the other hand, conceptually related analyses from the broader family of metrics for quantifying neural covariation have been developed in the context of other neuroimaging modalities (e.g., EEG, fNIRS); for example, correlated component analysis for EEG (Dmochowski et al., 2012, 2014), wavelet transform coherence for fNIRS (Cui et al., 2012; Dommer et al., 2012; Holper et al., 2012; Jiang et al., 2012, 2015; Nozawa et al., 2016; Hu et al., 2017), and adaptations of phase synchrony for fMRI (Glerean et al., 2012; Nummenmaa et al., 2014a, 2014b).

## Conclusion

With social and affective neuroscience aiming to study brain processes involved in rich and naturalistic situations, ISC analysis adds a valuable tool to our methodological arsenal. At base, this tool enables us to filter out idiosyncratic signals and localize brain regions that encode stimulus qualities consistently across individuals without an explicit model of the stimulus. Recent extensions of this approach incorporate spatially distributed response patterns and measure functional interactions between brain regions in real-life natural contexts. These tools not only provide a measure of the reliability of neural representation, but can provide a window into how humans, as social organisms, share and transmit information from person to person.

## Acknowledgements

We would like to thank Christopher J. Honey and Janice Chen for helpful comments on the manuscript, Qihong Lu, Manoj Kumar, Mai Nguyen, Meir Meshulam, Asieh Zadbood, and Lorenzo de Angelis for comments on the code and tutorial, and Jukka-Pekka Kauppi for helping with Appendix C.

## Funding

This work was supported by the National Institutes of Health (R01 MH112566-01 to U.H. and S.A.N.), by the Netherlands Organization for Scientific Research (VICI: 453-15-009 to C.K. and VIDI 452-14-015 to VG) and the European Research Council of the European Commission (ERC-StG-312511 to C.K.).

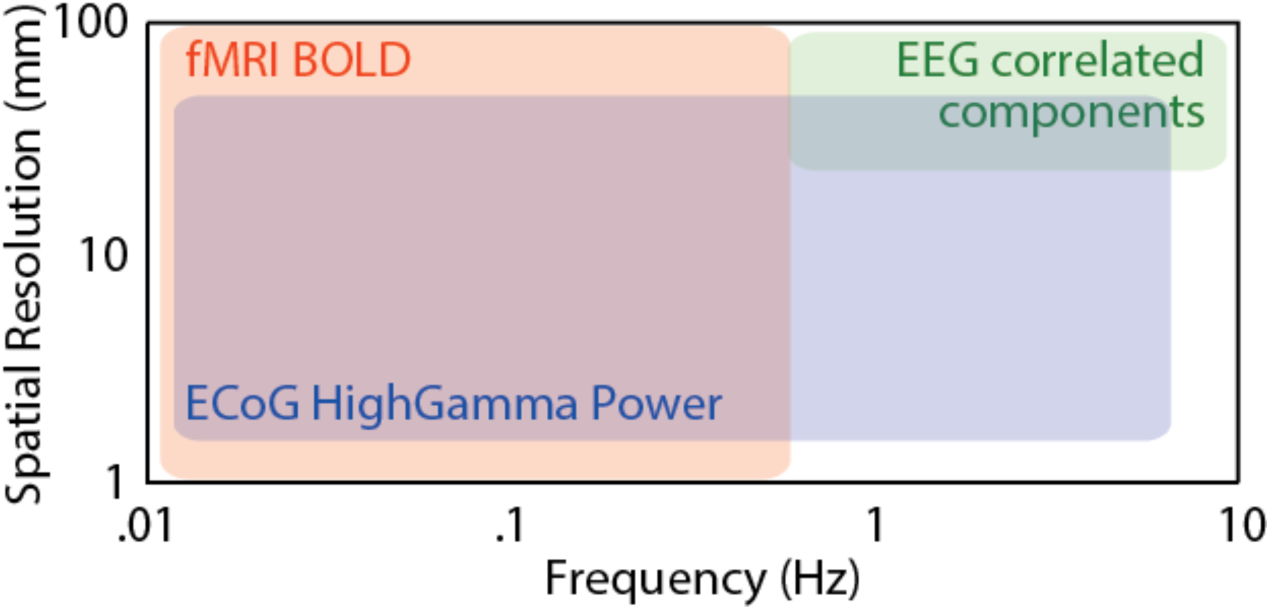

## Box 1. Spatiotemporal considerations

ISC can be evaluated both spatially and temporally at multiple scales. It is important to measure brain activity at a scale relevant to the phenomenon of interest.

### Temporal scale

Brain activity fluctuates at a number of frequencies that range from milliseconds (e.g., action potentials; >100 Hz) to the minute-long fluctuations often seen in BOLD activity (0.01 Hz). Our cognition also operates over multiple temporal scales: our visual system is able to perceive flicker at up to 10 Hz, while our emotions fluctuate at frequencies typically below 0.1 Hz. To use ISC, we must determine the frequency range relevant to our function of interest and adopt a measurement technique that is sensitive to fluctuations of brain activity in that frequency band (see above). Band-pass filtering brain activity measurements to that frequency band can help zoom in on a function of interest. Multi-modal approaches combining fMRI with high-temporal-resolution technologies such as EEG and ECoG can provide insights as to what frequency bands contribute most to ISCs (Mukamel et al., 2008; Liu et al., 2017; Haufe et al., 2018).

### Spatial scale

The brain is organized at multiple spatial scales and different functional topographies are multiplexed on the cortical sheet. For example, in V1, orientation is represented at the sub-millimeter scale, while eccentricity is represented at the centimeter scale; downstream visual areas encode overlapping representations of eccentricity, object category, and other object properties. Importantly, spatial (and spatiotemporal) ISC is sensitive to shared representations that are encoded across distributed response patterns. Investigating fine-grained representations across subjects requires a fine spatial scale of measurement, and responses may be consistent across subjects at some spatial scales and not others. At the centimeter scale, representational maps are relatively consistent across subjects, providing a rationale for applying ISC analyses to smoothed fMRI data (see Appendix A). To study ISCs at a finer scale, anatomical alignment will not sufficiently align functional topographies across individuals; it will be essential to functionally realign voxels from different brain using independent data prior to computing ISCs in a shared representational space (Haxby et al., 2011; Guntupalli et al., 2016).

### Appendix A: Typical preprocessing pipeline

In the following, we recommend a basic preprocessing pipeline intended to precede ISC analysis, followed by an example of how to implement this in SPM (Friston et al., 2007); however, several other commonly used software tools implement components and variations of this preprocessing pipeline, including AFNI (Cox, 1996), fMRIPrep (Esteban et al., 2018), and FSL (Smith et al., 2004). When acquiring data, we recommend locking the stimulus onset to the scanner triggers (indicating TR onsets) and logging all triggers. We also recommend using ascending or descending slice acquisition on the scanner rather than interleaved sequences. Slice timing correction may be appropriate, but is not universally recommended. Rigid-body registration is then used to correct for head motion over the course of the scan. Note that motion correction does not fully ameliorate the negative effects of head motion. Spatial normalization is then used to transform each subject’s data into a common coordinate space. We recommend using either volumetric nonlinear anatomical normalization to a high-resolution template in MNI space (e.g., Fonov et al., 2011) or surface-based normalization based on sulcal curvature (Fischl et al., 1999). Typically, each subject’s functional data are first aligned to that subject’s high-resolution anatomical image using an affine transformation, and the anatomical image is normalized to MNI space. These transformations should be concatenated and applied in a single step to avoid resampling and interpolating the data multiple times. For conventional temporal ISC analyses, we recommend spatially smoothing the data using a 4–8 mm full width at half maximum (FWHM) Gaussian kernel (Pajula and Tohka, 2014). For spatial and spatiotemporal ISC analysis, we recommend limited (e.g., 4 mm FWHM) or no smoothing at all. Although ISCs filter out idiosyncratic noise signals to some extent, we nonetheless recommend filtering out nuisance variables prior to ISC analysis. Commonly-used nuisance variables include head motion, principal component time series from anatomically segmented cerebrospinal fluid masks (Behzadi et al., 2009), framewise displacement (Power et al., 2013), linear and quadratic trends, as well as sine/cosine bases comprising a high-pass (e.g., 0.007 Hz) or band-pass (e.g., 0.01–0.1 Hz) filter. Nuisance variables and temporal filters should be combined into a single regression model to ensure that noise signals are not injected back into the data and to properly estimate degrees of freedom (Lindquist et al., 2019). If performing a volumetric analysis, ISCs should be computed within a brain or gray matter mask to reduce the number of subsequent tests (surface-based analyses preclude this step).

*Workflow* (in SPM terminology as an example)

a) Data Acquisition with conditions synchronized to TR and ascending or descending slice acquisition using the 4D NIfTI format.

b) Apply Slice timing correction to all images.

c) Apply Realign (Estimate and Reslice), with options: Register to mean and Resliced images: Mean Image Only.

d) Use Coregister Estimate, set Reference Image to the mean EPI image from (c) and Source Image to the anatomical T1 image.

e) Use Segment, Volumes: the coregistered T1 image from (d). Then select Deformation Fields: Inverse+Forward to save the functions that will transform the T1 and the EPI images to MNI space.

f) Normalize Write: Select the deformation field from step (e), and select all the EPI volumes and the T1 image and the Gray-Matter segment.

h) The normalized EPI images should then be smoothed using a FWHM kernel of 2–2.5 times the original voxel size (e.g., at 8 mm FWHM for data acquired at 3 × 3 × 3 mm; Pajula and Tohka 2014).

f) An average Gray-Matter should be created, by selecting all the normalized Gray-Matter segmentations of all the subjects. For that use ImCalc. Set Data Matrix to yes, select all your normalized gray matter images (N = number of subjects), and use Expression: mean(X).

g) The average Gray-Matter segmentation should be thresholded to provide a gray matter mask to use for ISC analyses. A threshold of around 0.25 is often useful. To do this use ImCalc, select the average normalized Gray, and set Expression to i1 > 0.25.

i) Optional: regress out the motion parameters and the average signal in the white matter and CSF, and apply a high-pass filter (100 s; Stephens et al., 2013). The simplest way to do this is to extract the signal from the segmentations using the normalized CSF and White-Matter of the participant (e.g., using MarsBaR), and then build a first-level model with co-variates only, including the motion parameters, the extracted average time-course of CSF and White-Matter from MarsBaR, and a 100-second high-pass filter. Then evaluate the model selecting the option ‘save residuals’. These residuals are then the signals that are used for further analyses.

j) The relevant segments of each subject’s data (i.e., the residuals from i) are then trimmed (to remove the first 6–10 seconds of each epoch capturing nonspecific stimulus onset) and *z*-scored (standardized to zero mean and unit variance) for each voxel and segment, then concatenated into a single 4D NIfTI file. If you want to compare multiple conditions, you can create one 4D NIfTI file for each condition. You may then mask the functional data using a gray-matter segmentation mask.

### Appendix B: Parametric paired test for two conditions

a) For each participant, you should have one 4D NIfTI file for each condition in a standard space; e.g., `sub-01_task-intact_bold.nii.gz`, `sub-01_task-scrambled_bold.nii.gz`, `sub-02_task-intact_bold.nii.gz`, `sub-02_task-scrambled_bold.nii.gz`, etc. We’ll compute leave-one-out ISCs separately for each condition.

b) We provide two simple scripts to compute the leave-one-out ISCs. For those familiar with MATLAB, use the `isc_loo.m` MATLAB script (specify the input and output directories). We also provide a Python command-line interface (CLI) called `isc_cli.py` for Linux or Mac (requires an installation of Python 3 with the NumPy/SciPy and NiBabel modules). You can run the program on the command line using `python3 isc_cli.py`; alternatively, you can make the script executable by running `chmod +x isc_cli.py`, after which you can run the program using `./isc_cli.py`. The `isc_cli.py` program requires input (`-i` or `--input`) and output (`-o` or `--output`) specifying filenames, and accepts a handful of optional arguments for specifying a mask filename (`-m` or `--mask`), *z*-scoring input time series (`-z` or `--zscore`), Fisher *z*-transforming output ISCs (`-f` or `--fisherz`), or computing a summary statistic (`-s` or `--summarize`; either `mean` or `median`). Usage example:

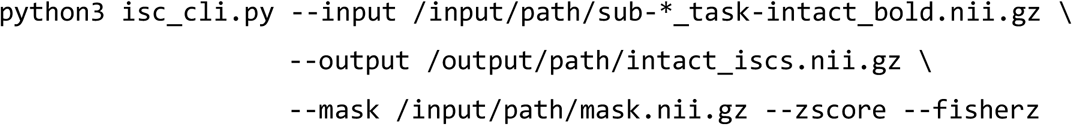

c) To test for a statistically significant difference between the Fisher *z*-transformed ISC values for each condition, perform a group-level analysis using standard fMRI analysis software. For example, in AFNI, use `3dttest++` with the `-setA`, `-setB`, and `-paired` options. In SPM, this entails using ‘specify 2^nd^-level’ and a paired *t*-test to compare two conditions. With large sample sizes, you may also opt to correlate the ISC values with subject-specific behavioral or demographic variables (e.g., how well each participant comprehended a story stimulus).

e) Correct the resulting *p*-values for multiple tests by, e.g., controlling FDR at .05 (for example, by submitting the *p*-values to AFNI’s `3dFDR`).

f) Visualize the resulting ISC maps by plotting any ISC values surpassing FDR controlled at .05. Typically plotting the ISC values themselves (or the difference in ISC values across conditions) after thresholding by the significance level is preferred to plotting *p*-values directly, as the ISC values indicate the effect size (G. Chen et al., 2017a). As correlation values range from –1 to +1, it may be appropriate to use a symmetric divergent (bipolar) colormap; however, ISC values are typically positive in practice, meaning a sequential (unipolar) colormap may be appropriate as well. We recommend using perceptually uniform colormaps and avoiding perceptually non-uniform colormaps (such as ‘jet’).

### Appendix C: Pairwise ISC analysis using the ISC Toolbox

*a) How to install the ISC Toolbox (Kauppi et al., 2014)*

Go to the Source Code https://www.nitrc.org/scm/?group_id=947 and click on “Download The Nightly SVN Tree Snapshot”.

Unzip the zip file in directory of choice.

Follow the instructions in readme.txt to install the atlas files from FSL.

*b) How to perform an analysis using the ISC Toolbox*

This will perform a pairwise correlation analysis for a single condition (i.e., *r* > 0).

1. Start MATLAB and make the directory in which you installed the toolbox in step A your current directory.
2. Launch setISCToolboxPath to add the different tools to your MATLAB path.
3. Start the analysis toolbox by typing ISCanalysis. This will open a GUI.

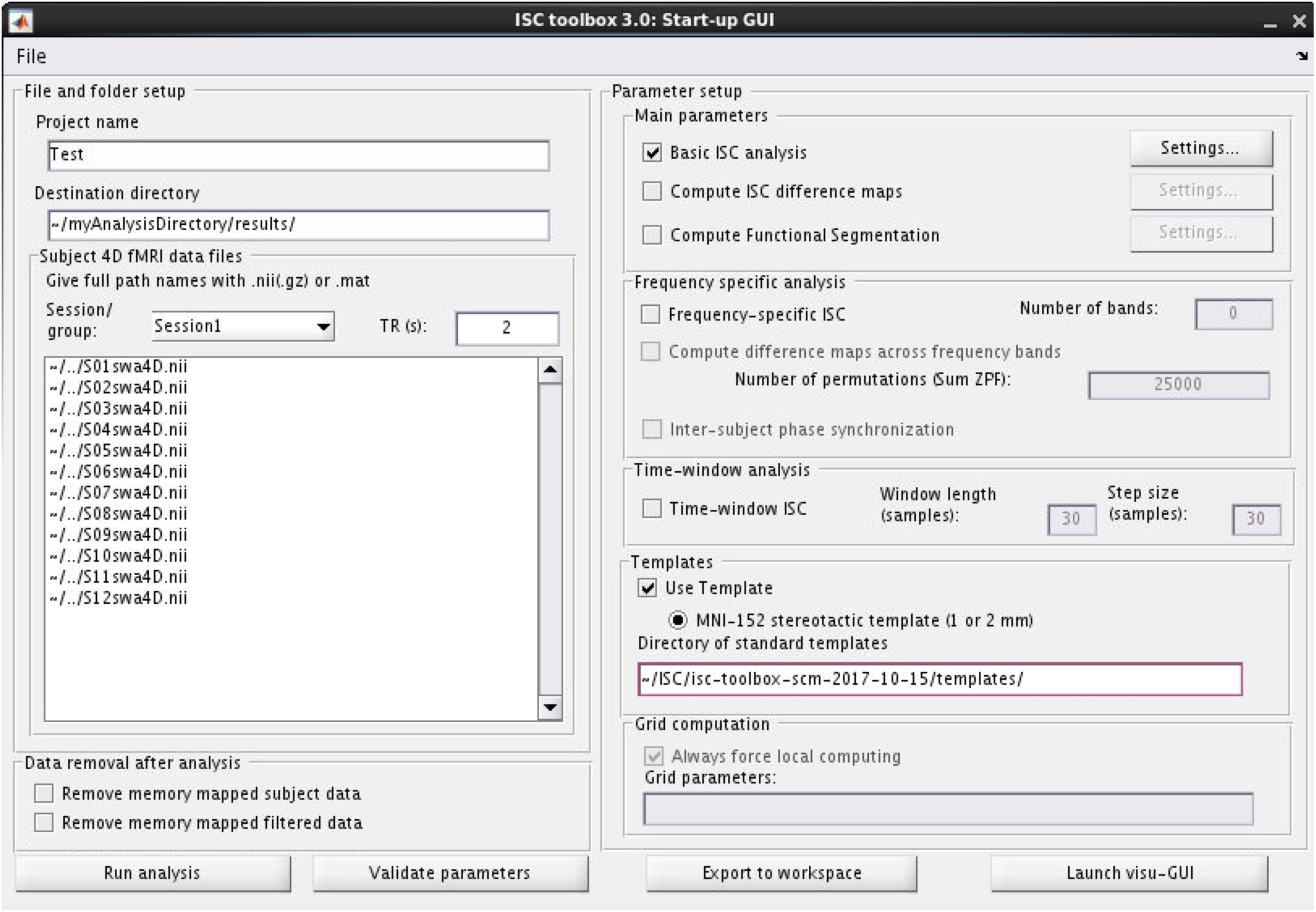 Screenshot from the GUI of the ISC Toolbox. File names are abbreviated here for illustration purposes.
4. Give the project a name.
5. Indicate the Destination Directory into which results will be copied.
6. Select the 4D NIfTI files that contain the preprocessed, clipped, concatenated data from each subject. These files must all have exactly the same number of volumes. You must copy the full path of each of these files into the window.
7. Specify your TR in seconds.
8. Select “Basic ISC analysis” in main parameters, and go to Settings… and select Calculate average ISC maps.
9. Set the number of realizations to 1,000,000.
10. Do not select frequency specific analyses or time window analyses.
11. Then press “Validate parameters”.
12. You should see a message confirming validation in the MATLAB kernel.
13. Press “Run analysis”.
14. This will take a while and will create a directory with results where you selected your destination directory.
15. In particular, in the subdirectory results, you will find two useful files: ISCcorrmapBand0Session1.nii and Session1Band0ThresholdsWin0.csv. The former contains the average pairwise ISC for each voxel, the latter the values at which to threshold this map to respect a certain threshold, in the following order:

% 1. *p* < 0.05, no multiple comparisons correction

% 2. *p* < 0.05, FDR corrected using independence or positive dependence assumption

% 3. *p* < 0.05, FDR corrected (no assumptions)

% 4. *p* < 0.05, Bonferroni corrected

% 5. *p* < 0.005, no multiple comparisons correction

% 6. *p* < 0.005, FDR corrected using independence or positive dependence assumption

% 7. *p* < 0.005, FDR corrected (no assumptions)

% 8. *p* < 0.005, Bonferroni corrected

% 9. *p* < 0.001, no multiple comparisons correction

% 10. *p* < 0.001, FDR corrected using independence or positive dependence assumption

% 11. *p* < 0.001, FDR corrected (no assumptions)

% 12. *p* < 0.001, Bonferroni corrected

You can then load this map in your preferred viewer (e.g., SPM with the anatomy toolbox). A table of significant clusters can be easily exported into a text file.

### Appendix D: ISC analysis using BrainIAK in Python

This manuscript is accompanied by a GitHub repository: https://github.com/snastase/isc-tutorial. This repository contains a Jupyter Notebook tutorial that introduces basic ISC analyses and statistical tests implemented in Python using BrainIAK. The code can be interactively executed and modified in the cloud using the free Google Colaboratory notebook environment. To use the tutorial notebook interactively, click *Open in playground*; this will allow you to edit and run the code cells (you may need to log into a Google account). Use *File* > *Save a copy in Drive*… or *Save a copy in GitHub*… to save your changes. We recommend exploring the tutorial notebook via the browser-based Colaboratory environment, you may also download the notebook file and run it locally using Jupyter Notebook (with some minor modifications; see below). In addition to the tutorial notebook, we include some convenience functions: an example MATLAB script (`isc_loo.m`) and a simple, Python-based command-line program (`isc_cli.py`) for computing leave-one-out ISCs.

#### Installing Python

To install Python, we recommend using the Anaconda distribution: https://www.anaconda.com/distribution. Select the installer compatible with your operating system and click on the *Download* button corresponding to Python 3 (e.g., Python 3.7). When the download is complete, open the installer and follow the installation instructions. In the tutorial notebook, we install some nonstandard Python libraries not included in the default Anaconda distribution, namely NiBabel (https://nipy.org/nibabel), Nilearn (https://nilearn.github.io), and BrainIAK (https://brainiak.org). These can be installed from the command line using, e.g., `pip install nibabel`. BrainIAK is supported for Linux and MacOS and requires an installation of Python version 3.4 or higher. Follow the instructions at the following link to install BrainIAK and its dependencies using `pip` or `conda`: http://brainiak.org/docs/installation.html. If you are unable to install BrainIAK, you can use a demo version of the ISC functions by downloading the `isc_standalone.py` module from the GitHub repository.

#### Tutorial software

To use the tutorial software locally, you can either (*a*) clone the GitHub repository to a local directory on the command line using git clone https://github.com/snastase/isc-tutorial.git; or (*b*) click on the green *Clone or download* button, click *Download ZIP*, and extract the contents of the archive in a local directory. To open the tutorial notebook, navigate into the `isc-tutorial` directory (or `isc-tutorial-master` if you downloaded the ZIP archive) and launch Jupyter Notebook (e.g., by running `jupyter notebook` from the command line). This will open a browser window displaying the files in the directory where you launched Jupyter Notebook. Click on the `isc_tutorial.ipynb` notebook file, which will open another browser window containing the interactive tutorial notebook. The tutorial notebook contains both explanatory text and code cells. The first code cell in the tutorial notebook is intended to install the software requirements for BrainIAK in the Linux cloud instance hosted by Google Colab and should not be executed if running the notebook locally. If you have already successfully installed BrainIAK, you can skip to the second code cell to import the necessary BrainIAK functions. The third code cell downloads the `isc_standalone.py` file for those unable to install BrainIAK, and will not be necessary if you cloned the GitHub repository (which already contains a copy of the `isc_standalone.py` file). For those working locally without a BrainIAK installation, skip to the fourth cell which imports the necessary functions from the `isc_standalone.py` module. To execute a code cell, click on the cell, then either click the *Run* button (or the run arrow to the left of the cell in Google Colab) or type *Shift* + *Enter*. Click on a code cell to edit it. Note that the *Open in Colab* button at the top of the notebook will open the notebook in a Google Colab cloud instance as described above.

#### ISC analysis

The `isc` function in BrainIAK takes in NumPy arrays comprising BOLD time series for one or more voxels or ROIs across two or more subjects and returns ISC values for each voxel or ROI. The `pairwise` argument can be used to toggle between the pairwise approach (`pairwise=True`) and the leave-one-out approach (`pairwise=False`). By default, this function returns ISC values for either all pairs of subjects or each left-out subject; however, you can supply a `summary_statistic` (‘mean’ or ‘median’), which will yield a single summary ISC statistic across pairs or left-out subjects. If mean ISCs are requested, Fisher *z*-transformation is applied appropriately.

#### Statistical tests

BrainIAK currently supplies four nonparametric methods for statistically evaluating ISCs. The `timeshift_isc` and `phaseshift_isc` functions operate directly on response time series, applying circular time-shift or phase randomization prior to re-computing ISC at each iteration of the resampling test. On the other hand, the `bootstrap_isc` and `permutation_isc` functions operate on ISC values, applying a bootstrap hypothesis test and permutation tests, respectively. The `permutation_isc` function can be provided `group_assignment` labels to perform a two-sample test, or performs a one-sample test using a sign-flipping procedure. Similar to the core `isc` function, these nonparametric tests can be supplied with `pairwise` and `summary_statistic` arguments. Each statistical test returns the observed ISC, *p*-values based on the null distribution, and the null distribution itself (as well as confidence intervals in the bootstrap hypothesis test).

#### ISFC analysis

To perform ISFC analysis, supply BOLD time series data for two or more voxels (or ROIs) across two or more subjects to the `isfc` function. ISFCs can be computed using either the pairwise (`pairwise=True`) or leave-one-out (`pairwise=False`) approaches. The `isfc` function returns a tuple containing a condensed vector of off-diagonal ISFC values and the diagonal ISC values (`vectorize_isfcs=True`), or a 3-dimensional NumPy array where the first dimension corresponds to pairs or left-out subjects and the latter two dimensions correspond to the voxel-by-voxel (redundant) ISFC matrix (`vectorize_isfcs=False`). If a summary statistic is supplied, only the voxel-by-voxel ISFC matrix is returned, collapsing across pairs or left-out subjects.

